# Cryo-EM structure and kinetics reveal electron transfer by 2D diffusion of cytochrome *c* in the yeast III-IV respiratory supercomplex

**DOI:** 10.1101/2020.11.27.401935

**Authors:** Agnes Moe, Justin Di Trani, John L. Rubinstein, Peter Brzezinski

**Author notes:** A.M. and J.D.T. contributed equally to this work.

## Abstract

Energy conversion in aerobic organisms involves an electron current from low-potential donors, such as NADH and succinate, to dioxygen through the membrane-bound respiratory chain. Electron transfer is coupled to transmembrane proton transport that maintains the electrochemical proton gradient used to produce ATP and drive other cellular processes. Electrons are transferred between respiratory complexes III and IV (CIII and CIV) by water-soluble cyt. *c*. In *S. cerevisiae* and some other organisms, these complexes assemble into larger CIII_2_CIV_1/2_ supercomplexes, the functional significance of which has remained enigmatic. In this work, we measured the kinetics of the *S. cerevisiae* supercomplex’s cyt.*c*-mediated QH_2_:O_2_ oxidoreductase activity under various conditions. The data indicate that the electronic link between CIII and CIV is confined to the surface of the supercomplex. Cryo-EM structures of the supercomplex with cyt. *c* reveal distinct states where the positively-charged cyt. *c* is bound either to CIII or CIV, or resides at intermediate positions. Collectively, the structural and kinetic data indicate that cyt. *c* travels along a negatively-charged surface patch of the supercomplex. Thus, rather than enhancing electron-transfer rates by decreasing the distance cyt. *c* must diffuse in 3D, formation of the CIII_2_CIV_1/2_ supercomplex facilitates electron transfer by 2D diffusion of cyt. *c*. This mechanism enables the CIII_2_CIV_1/2_ supercomplex to increase QH_2_:O_2_ oxidoreductase activity and suggests a possible regulatory role for supercomplex formation in the respiratory chain.

**Significance Statement:** In the last steps of food oxidation in living organisms, electrons are transferred to oxygen through the membrane-bound respiratory chain. This electron transfer is mediated by mobile carriers such as membrane-bound quinone and water-soluble cyt. *c*. The latter transfers electrons from respiratory complex III to IV. In yeast these complexes assemble into III_2_IV_1/2_ supercomplexes, but their role has remained enigmatic. This study establishes a functional role for this supramolecular assembly in the mitochondrial membrane. We used cryo-EM and kinetic studies to show that cyt. *c* shuttles electrons by sliding along the surface of III_2_IV_1/2_ (2D diffusion). The structural arrangement into III_2_IV_1/2_ supercomplexes suggests a mechanism to regulate cellular respiration.

## Introduction

Aerobic organisms obtain energy by linking oxidation of food to the synthesis of adenosine triphosphate (ATP). An intermediate step in this process is translocation of protons across a membrane by a series of integral membrane proteins collectively known as the respiratory chain. In eukaryotes, the respiratory chain is found in the inner mitochondrial membrane. Transmembrane proton translocation is driven by electron transfer from NADH and succinate to O_2_, which renders the mitochondrial matrix more negative (*n* side) and the intermembrane space more positive (*p* side). The resulting electrochemical proton gradient is used for ATP production by ATP synthase and for transmembrane transport processes.

In *Saccharomyces cerevisiae*, NADH donates electrons to dehydrogenases Nde1, Nde2, and Ndi1 (1), which reduce membrane-bound quinone (Q) to quinol (QH_2_). Succinate dehydrogenase also contributes to the pool of reduced QH_2_ in the membrane. Quinol diffuses in the hydrophobic core of the lipid bilayer to donate electrons to cytochrome *c* reductase (also known as cyt. *bc*_1_ or complex III), which forms an obligate homodimer (CIII_2_) (for reviews, see (2–5)). Within CIII, electrons are transferred from QH_2_ to cyt. *c*_1_, which is a component of CIII, in a series of reactions known as the proton-motive Q-cycle (see **Fig. 1A** and (3, 5)). Briefly, binding of QH_2_ in a site near the *p* side, called Q_P_ (or Q_o_) is followed by transfer of one of its electron, first to an FeS center and then to cyt. *c*_1_. This oxidation of QH_2_ is linked to release of its two protons to the *p* side of the membrane. The second electron from the resulting semiquinone (SQ^•-^) in the Q_P_ site is transferred via hemes *b*_L_ and *b*_H_ to bound Q in the Q_N_ (Q_i_) site, reducing it to SQ^•-^. Repetition of the same sequence of events leads to reduction of the SQ^•-^ in the Q_N_ site to QH_2_, which abstracts two protons from the *n* side of the membrane and subsequently equilibrates with the reduced QH_2_ pool.

**Figure 1.**
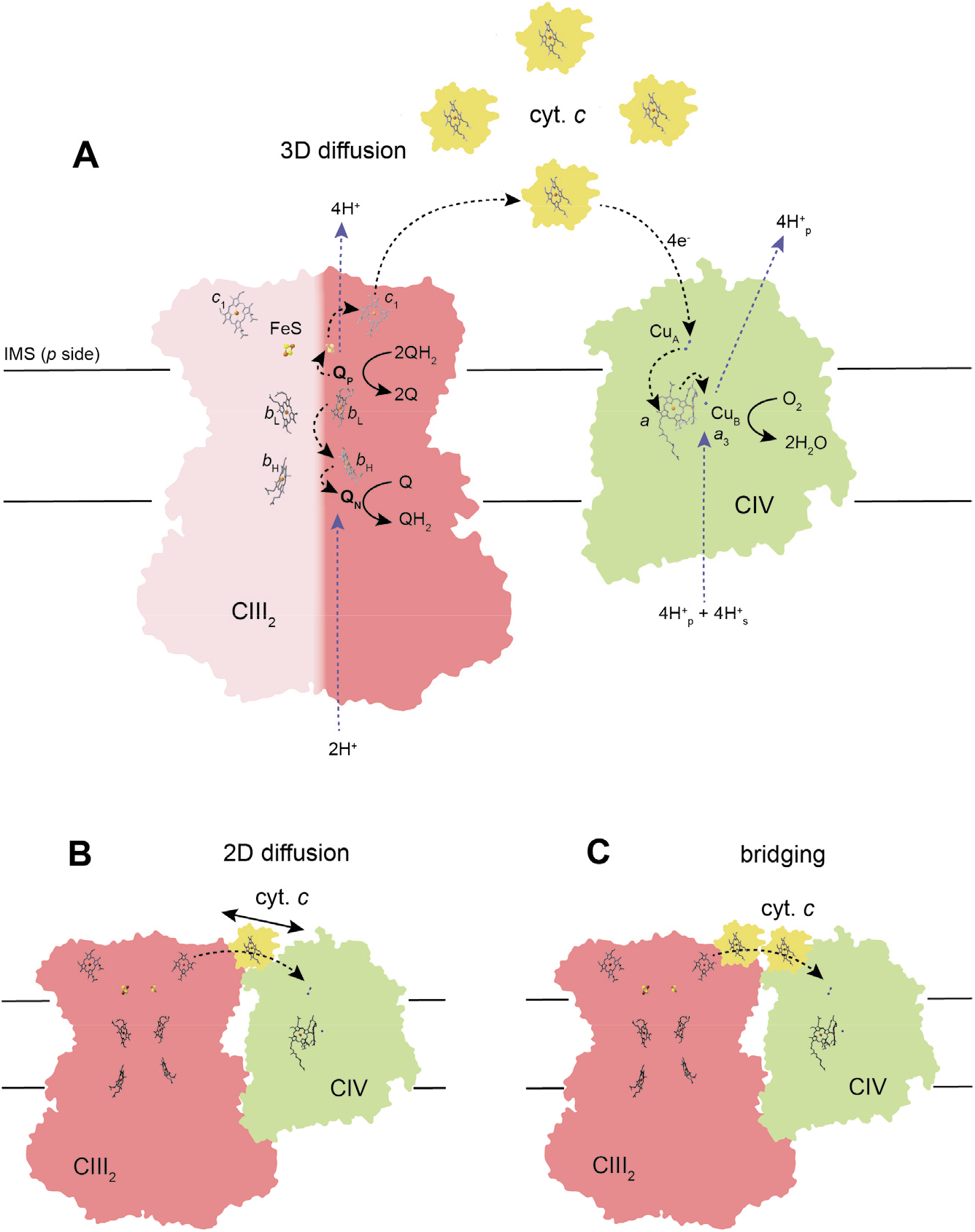
Reactions catalyzed by the CIII_2_ and CIV, and possible models for electron transfer between them. Q_N_ and Q_P_ indicate the two quinol/quinone-binding sites. The black and blue arrows indicate electron-transfer and proton-transfer reactions, respectively. In CIV, H^+^_p_ and H^+^_s_ indicate protons that are pumped or used as substrate for reduction of O_2_ to H_2_O, respectively. Only reactions in one half of the CIII_2_ dimer are indicated. These reactions occur independently in the two halves of the dimer. Transfer of electrons between CIII_2_ and CIV via the 3D diffusion of cyt. *c* is depicted. (**B**) Electron transfer from CIII_2_ to CIV via 2D diffusion of a cyt. *c* associated with the CIII_2_CIV_2_ supercomplex. (**C**) Electron transfer from CIII_2_ to CIV via a bridge formed by more than one bound cyt. *c*.

Electrons from cyt. *c*_1_ within CIII are transferred to the water-soluble mobile one-electron carrier cyt. *c*, which resides in the mitochondrial intermembrane space (**Fig. 1A**). Reduced cyt. *c* binds to the *p*-side surface of cytochrome *c* oxidase (also known as cyt. *aa3* or complex IV) where its electron is transferred first to CIV’s di-nuclear copper site, Cu_A_, then heme *a*, and then the binuclear heme *a*_3_ and Cu_B_ catalytic site. Transfer of a second electron from a second cyt. *c* to CIV leads to reduction of both heme *a*_3_ and Cu_B_, allowing O_2_ to bind to the heme *a*_3_ iron. This O_2_ is reduced to water after transfer of a total of four electrons from four cyt. *c* molecules to the catalytic site. Each electron transfer to the catalytic site is linked to uptake of one proton from the *n* side of the membrane and pumping of one proton across the membrane from the *n* to the *p* side (**Fig. 1A**) (for review, see (6, 7)).

Each respiratory complex can function independently of the others and early studies supported a model where the complexes diffuse independently in the mitochondrial inner membrane (8) (**Fig. 1A**). However, more recently a variable fraction of the respiratory complexes have been shown to form larger supercomplexes consisting of two or more components of the respiratory chain (9–13). Supercomplexes can be isolated with preserved enzymatic activity (14, 15) and the overall arrangement of complexes within supercomplexes has been determined in cells from mammals, yeast, and plants (16). In addition, respiratory supercomplexes with different compositions and stoichiometries of components have been isolated using mild detergents and their high-resolution structures have been determined by single particle electron cryomicroscopy (cryo-EM) (reviewed in (17)).

In *S. cerevisiae* essentially all CIVs are part of supercomplexes (18) composed of CIII_2_ flanked by either one or two copies of CIV (9, 10, 18–26). Recent studies showed only minor structural changes in the individual complexes upon association (27–29), which suggests that CIII-CIV binding does not result in functional differences in the components and that supercomplex formation alters only their proximity.

Electron transfer from CIII to CIV within the supercomplex requires cyt. *c* (28). The cyt. *c* docking sites in CIII and CIV are of 60 to 70 Å apart, which is too far to allow direct electron transfer through a single stationary cyt. *c* on the supercomplex surface. Thus, electron transfer between the two complexes must occur via one of three scenarios (**Fig. 1A-C**): (*i*) three-dimensional (3D) diffusion of cyt. *c* between CIII and CIV (8), (*ii*) lateral diffusion of cyt. *c* along the supercomplex surface (26, 30–33) (see also (34)), or (*iii*) two or more cyt. *c* molecules that bind simultaneously on the supercomplex surface to bridge CIII and CIV. Here, we explored these possibilities to address the functional significance of supercomplex formation with combined structural and functional studies of the *S. cerevisiae* supercomplex with added cyt. *c*. The kinetic data exclude the possibility that electron transfer between CIII and CIV by cyt. *c* involves 3D diffusion of reduced cyt. *c* between its CIII and CIV binding sites. The structural data show an ensemble of states where cyt. *c* is bound either to CIII, CIV, or at intermediate positions between the two on the supercomplex surface. Collectively, these results indicate that electron transfer between CIII and CIV occurs along the surface of the supercomplex, suggesting a mechanism to regulate the redox state of the cyt. *c* pool by altering the CIII:CIV ratio and through association/dissociation of supercomplexes upon changing environmental conditions.

## Results

### Isolation of the yeast CIII_2_CIV_1/2_ supercomplex

The *S. cerevisiae* CIII_2_CIV_1/2_ supercomplex was purified using a FLAG tag on the Cox6 subunit of CIV (35) (**Figs. S1AB**). Absorption spectroscopy of the preparation (**Fig. S1C**) gave difference spectra similar to previous results (28) and yielded a heme *a*:*b*:*c* ratio of ~1.7:2:1, which is consistent with a mixture of CIII_2_CIV_1_ and CIII_2_CIV_2_ supercomplexes, as observed previously (27–29). Furthermore, the heme *b*:*c* ratio of ~2:1 indicates that the only heme *c* present was from the bound cyt. *c*_1_ heme of CIII and not from co-purifying soluble cyt. *c*. Association of soluble cyt. *c* with CIII and CIV is known to depend on the ionic strength of the surrounding solution (36). Consequently, all subsequent experiments were done in buffer with 150 mM KCl at near-physiological ionic strength.

### Activity of CIII and CIV

To probe the catalytic activity of CIII_2_ and CIV within the CIII_2_CIV_1/2_ supercomplex, the activity of each component was measured separately. The supercomplex was maintained in solution by the mild detergent glycodiosgenin (GDN). The activity of CIII_2_ was measured at 90 ± 20 e^-^/s by following reduction of cyt. *c* by decylubiquinol (DQH_2_) spectrophotometrically while CIV was inhibited by cyanide. The activity of CIV was measured at 450 ± 20 e^-^/s using a Clarke electrode to follow the reduction of oxygen by cyt. *c* that was maintained in a reduced state by ascorbate. The activities of free CIII_2_ and CIV were then measured after disruption of the supercomplex into CIII_2_ and CIV components by incubation with the detergent *n*-Dodecyl β-D-maltoside (DDM) (37). These activities were 60 ± 15 e^-^/s and 370 ± 30 e^-^/s, respectively, showing that the exchange of detergent and dissociation of the supercomplexes had only minor effects on this maximal activity of the individual complexes when cyt. *c* is at saturating concentrations (see below). The error estimated for all measurements is ±SD for three technical replicates from two independent preparations.

### Supercomplex activity

The QH_2_:O_2_ oxidoreductase activity of the supercomplex was measured by monitoring the O_2_-reduction rate upon addition of DQH_2_ in the presence of varying amounts of *S. cerevisiae* cyt. *c* (**Figs. 2A** and **S2**). As observed previously, the supercomplex is not able to reduce O_2_ with DQH_2_ in the absence of exogenous cyt. *c* (**Fig. 2A**)(28). Hence, even when CIII_2_ and CIV are held in close proximity within a supercomplex, electron transfer does not take place between the two components without cyt. *c*. At a cyt. *c*:supercomplex ratio of approximately one (~20 nM cyt. *c*), the turnover rate was 15 ± 1 e^-^/s (**Fig. 2B)**.

**Figure 2.**
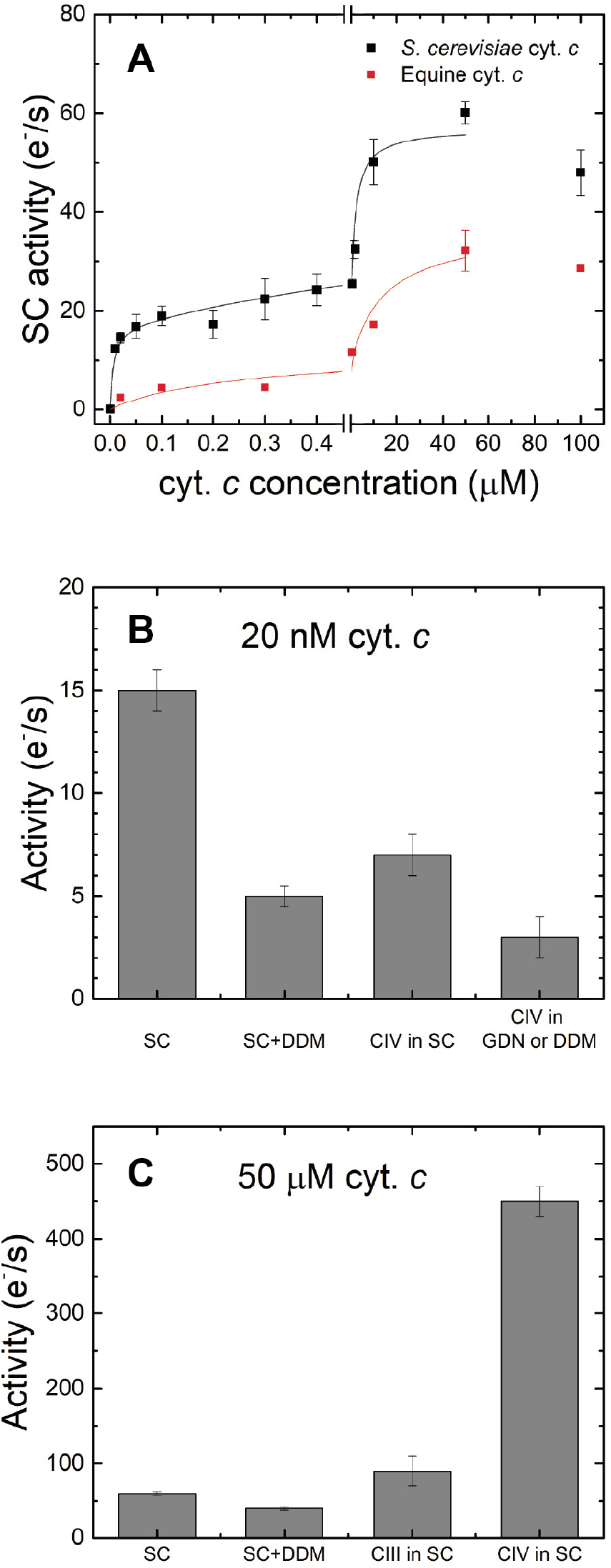
Catalytic activity. (**A**) The CIII_2_CIV_1/2_ supercomplex (SC) QH_2_:O_2_ oxidoreductase activity at different concentrations of *S. cerevisiae* or equine cyt. *c*. (**B**) The supercomplex QH_2_:O_2_ oxidoreductase activity at 20 nM cyt. *c* with intact (SC) and dissociated supercomplexes after incubation in DDM (SC+DDM). The cyt. *c* oxidation-O_2_ reduction activity was measured for CIV within a supercomplex (CIV in SC), for isolated CIV in GDN (from CIV peak in **Fig. S1A**), or after incubation in DDM (CIV in DDM). (**C**) The QH_2_:O_2_ oxidoreductase activity of the supercomplex at 50 μM cyt. *c* with intact (SC) and after dissociation of the supercomplex in DDM (SC+DDM). The maximum activities of CIII and CIV are also shown. Error bars are ±SD for three technical replicates from two independent preparations (except the 200-nM point in panel A, which is an average of a total of five replicates). For some points the error bars are smaller than the symbols. The solid lines were generated using Michaelis–Menten equations with *K*_m_ values 6 nM and 1.7 μM (yeast cyt. *c*) or 0.2 μM and 15 μM (horse cyt. *c*).

As the cyt. *c*:supercomplex ratio is increased gradually from 1 to 10 (200 nM cyt. *c*) the turnover rate increases only slightly from 15 to ~20 e^-^/s. However, as the cyt. *c*:supercomplex ratio is increased beyond 10, turnover increases more quickly, reaching a maximum value of ~60 e^-^/s at a cyt. *c*:supercomplex ratio of 2500 (50 μM cyt. *c*). This maximal rate is only slightly lower than the maximum turnover rate of CIII_2_ alone (~90 e^-^/s), but is much lower than the rate of CIV (~450 e^-^/s), suggesting that supercomplex activity is limited by the turnover rate of CIII_2_ (**Fig. 2C**). The rate decreases slightly at a cyt. *c*:supercomplex ratio of 5000 (100 μM cyt. *c*), the highest ratio used (**Fig. 2A**). This decrease may be due to cyt. *c* binding a “nonproductive” site at CIV (38) that interferes with cyt. *c* diffusion between the productive binding sites at CIII and CIV.

The low and high cyt. *c*:supercomplex regimes for the cyt. *c* titration in **Fig. 2A** yield apparent *K*_M_ values of ≤6 nM and ~1.7 μM, respectively. These values are distinct from the dissociation constant, *K*_d_, of cyt. *c* binding because in the QH_2_:O_2_ oxidoreductase experiment cyt. *c* mediates electron transfer between two functionally independent enzymes. Steadystate turnover of isolated *S. cerevisiae* CIV with isoform-1 cyt. *c* yielded biphasic Eadie-Hofstee plots with *K*_M_ values of ~100 nM and ~30 μM at low and high cyt. *c*:supercomplex ratios, respectively (39, 40). These *K*_M_ values are much larger than the apparent *K*_M_ values from **Fig. 2A** indicating that the QH_2_:O_2_ oxidoreductase activity of the supercomplex is not rate-limited by 3D diffusion of cyt. *c* to CIV.

Measurement of QH_2_:O_2_ oxidoreductase activity with non-physiological equine cyt. *c* yielded apparent *K*_M_ values of ~0.2 μM and ~15 μM (**Fig. 2A**), which is consistent with a weaker affinity of the mammalian cyt. *c* for the *S. cerevisiae* supercomplex. The larger of these apparent *K*_M_ values is similar to the ~14 μM *K*_M_ value obtained previously for equine cyt. *c* binding to *S. cerevisiae* CIV alone (40). Furthermore, with an equine cyt. *c*:supercomplex ratio of one, the QH_2_:O_2_ turnover rate for the intact supercomplex was only ~3 e^-^/s (**Fig. 2A**), which is similar to that of the disrupted supercomplex in DDM using *S. cerevisiae* cyt. *c* (**Fig. 2B**). These data suggest that with equine cyt. *c*, which binds the supercomplex weakly, activity is rate-limited by binding of cyt. *c* to CIV via 3D diffusion. The maximum supercomplex activity at an equine cyt. *c*:supercomplex ratio of 2500 was ~30 e^-^/s (**Fig. 2A**), consistent with an earlier study (28).

The lower apparent yeast cyt. *c K*_M_ values for the supercomplex (**Fig. 2A**) compared with CIV alone (see above), suggest that cyt. *c* binds more strongly to the supercomplex than to the independent CIV and CIII_2_. This difference in *K*_M_ is consistent with the supercomplex presenting a large binding surface for cyt. *c* between CIII_2_ and CIV. To test this hypothesis, we measured the QH_2_:O_2_ oxidoreductase activity of a supercomplex preparation after addition of DDM to disrupt the supercomplex into free CIII_2_ and CIV (37). With approximately one cyt. *c* per supercomplex (20 nM of each) the QH_2_:O_2_ oxidoreductase rate decreased from ~15 e^-^/s to ~5 e^-^/s upon disruption of the supercomplex (**Figs. 2** and **3**), showing that an increase in the average distance between CIII_2_ and CIV results in a decrease in the turnover rate. Indeed, the supercomplex’s QH_2_:O_2_ oxidoreductase activity of ~15 e^-^/s is higher than the cyt. *c*:O_2_ oxidoreductase of ~3 e^-^/s for CIV alone at the same cyt. *c*:CIV ratio. In measurements of the supercomplex activity cyt. *c* is only partially reduced, while in measurements of isolated CIV activity it is fully reduced. Consequently the 3 e^-^/s activity for CIV is an upper limit and the difference between supercomplex activity and CIV activity is even more pronounced than the experiment indicates. The activity of CIV was the same whether obtained from CIV in GDN (from the CIV peak shown in **Fig. S1A**) or from CIV in DDM (**Fig. 2B**), which shows that DDM does not interfere with cyt. *c* binding. The QH_2_:O_2_ oxidoreductase activity of dissociated supercomplex (~5 e^-^/s, SC+DDM) was slightly larger than the cyt. *c*:O_2_ oxidoreductase activity of CIV (~3 e^-^/s), presumably due to incomplete dissociation of supercomplexes upon addition of DDM. The cyt. *c*:O_2_ oxidoreductase activity of CIV within an intact supercomplex (~7 e^-^/s) was higher than that of isolated CIV (~3 e^-^/s) at a cyt. *c*:CIV ratio of one, which is consistent with the hypothesis that a large cyt. *c* binding surface in the supercomplex facilitates cyt. *c*. binding.

**Figure 3.**
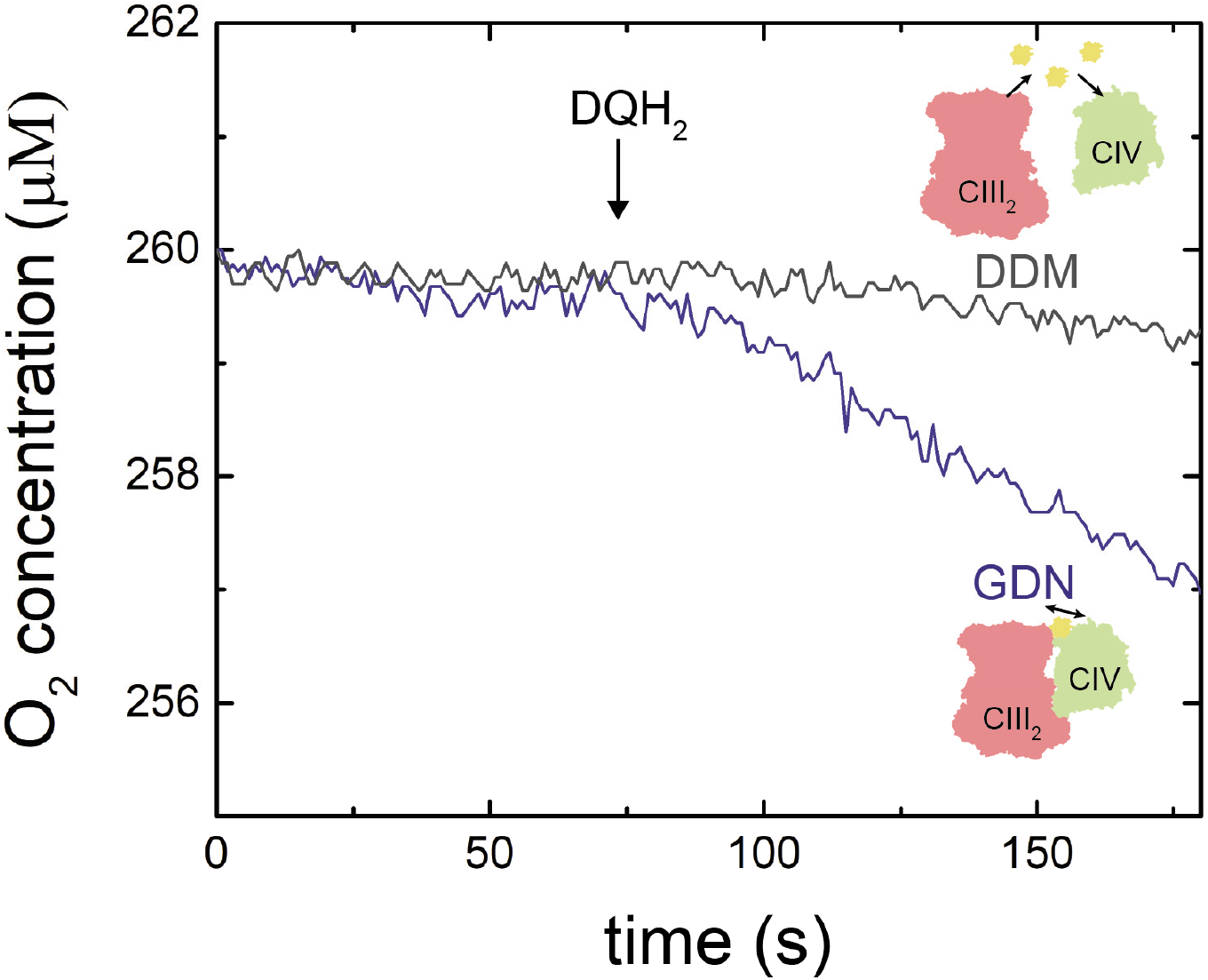
The QH_2_:O_2_ oxidoreductase activity of the supercomplex in GDN and DDM. The oxidoreductase rate was determined from the slope of the graph. Measurements were done with two different aliquots from the same supercomplex preparation in 0.01 % GDN.

These data indicate that cyt. *c* diffusion to CIV limits the QH_2_:O_2_ oxidoreductase activity for freely diffusing CIII_2_, CIV, and cyt. *c* at a cyt. *c*:supercomplex ratio of one. In contrast, with intact supercomplex, accumulation of cyt. *c* on the surface of the supercomplex facilitates the transfer of electron transfer between CIII and CIV. With a cyt. *c*:supercomplex ratio of 1 to 10 (20 to 200 nM cyt. *c*) this mechanism allows for a supercomplex activity of 15 to 20 e^-^/s (**Fig. 2A**).

### Structure determination by cryo-EM

In order to investigate the association of cyt. *c* with the surface of the supercomplex structurally, samples of the two were mixed in a ~1:12 ratio (11 μM supercomplex with 130 μM cyt. *c*) and the resulting preparation frozen on EM grids for analysis by cryo-EM (**Table S1** and **Fig. S3**). Cryo-EM resulted in 3D maps of supercomplexes consisting of both CIII_2_CIV_1_ and CIII_2_CIV_2_ (~2:1 CIII_2_CIV_1_:CIII_2_CIV_2_). As seen previously (27–29), in these structures the CIII_2_ dimer in the supercomplex is flanked by one or two CIV monomers (**Fig. S3B**). Focused refinement of the CIII_2_CIV_1_ structure with the entire dataset led to a map at 3.7 Å resolution (**Fig. 4Ai** and **S3C-E**). An atomic model of CIII_2_CIV_1_ from previous studies (27–29) fit well in the map (**Fig. 4Aii** and **S4A**). Local refinement of the CIII_2_ region led to small improvements in resolution for CIII_2_ while local refinement of the CIV region of led to a large improvement in resolution for CIV from ~5 Å to 4 Å resolution (**Fig. S5** and **S4BC**). No density was present for Rcf2, which was found in a recent CIII_2_CIV_1/2_ supercomplex structure (29).

**Figure 4.**
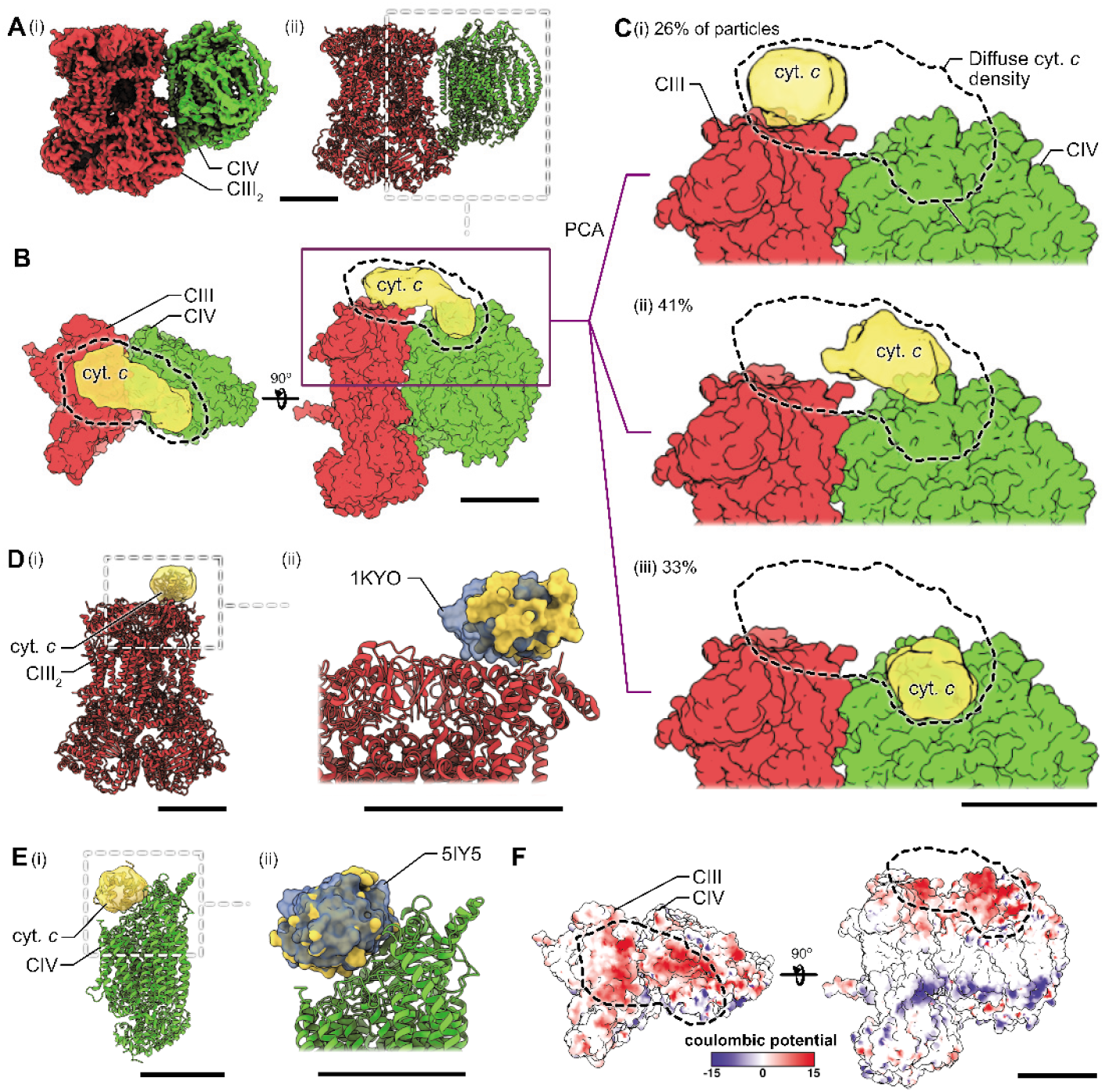
Cryo-EM analysis. (**A**) Cryo-EM map (***i***) and atomic model (***ii***) A of CIII_2_CIV_1_ supercomplex. (**B**) Surface representation of CIII_1_CIV_1_ supercomplex with density for cyt. *c* in yellow. (**C**) Particle images could be separated into populations that show cyt. *c* bound to CIII_2_ (***i***), in-between CIII_2_ and CIV (***ii***), and bound to CIV (***iii***). (**D**) An atomic model of yeast cyt. *c* (PDB: 1YCC (71)) (yellow) was fit into the structure from local refinement of the CIII_2_-cyt. *c* region of the map (***i***) and compared to the position of cyt. *c* from a previous structure of yeast CIII_2_-cyt. *c* (PDB: 1KYO (42) (blue) (***ii***). (**E**) The atomic model of yeast cyt. *c* (yellow) was fit into the structure from local refinement of the CIV-cyt. *c* region of the map (***i***) and compared to a crystal structure of bovine CIV-cyt. *c* (PDB: 5IY5 (43)) (blue) (***ii***). (**F**) Coulombic surface potential for supercomplex. Scale bars, 50 Å.

In the CIII_2_CIV_1_ map, an elongated density corresponding to cyt. *c* is found along the supercomplex surface, bridging the cyt. *c* binding sites in CIII and CIV (**Fig. 4B**). This density could arise due to multiple cyt. *c* molecules attached simultaneously to each supercomplex or incoherent averaging of a single cyt. *c* molecule with variable position along the supercomplex surface. To distinguish between these possibilities, the extended density was analyzed by principle component analysis, which is designed to detect continuous variability in protein structures (41) (**Fig. S6**). The analysis separated particle images into three clusters that were used to generate and refine three separate 3D maps. The three maps show the supercomplex with cyt. *c* bound at different positions (**Fig. 4C**). These structures indicate that the elongated density in the consensus structure in **Fig. 4B** is due to a single cyt. *c* with a position that varies continuously between CIII and CIV. In the maps cyt. *c* is seen bound at CIII (**Fig. 4Ci**), positioned between CIII and CIV (**Fig. 4Cii**), or bound at CIV (**Fig. 4Ciii**). The map with cyt. *c* bound to CIII was calculated from 26 % of particle images and focused refinement of the CIII_2_-cyt. *c* region shows clear density for a single well-defined cyt. *c* (**Fig. 4Ci, S7**). This map resembles a crystal structure of yeast CIII with cyt. *c* bound, (42) however, a rigid body fit of cyt. *c* from the crystal structure into the current map indicates a slight shift in its position (**Fig. 4D**).

The map showing cyt. *c* bound at CIV was obtained from 33 % of the particle images (**Fig. 4Ciii**). Similar to a previous cryo-EM study (29), focused refinement resulted in large improvements in resolution for the CIV region of the map, which allowed docking of a single cyt. *c* (**Fig. S8**) A recent crystal structure of mammalian CIV bound to cyt. *c* (43) could be fit into the map and closely matches the current model (**Fig. 4E**), indicating that the distance between cyt. *c* iron and Cu_A_ from CIV is similar in yeast and mammals.

The map with cyt. *c* approximately equidistant between CIII and CIV (**Fig. 4Cii**) was derived from 41 % of the particle images. In this intermediate position, the density for cyt. *c* remains blurred, with diffuse density extending all the way between the cyt. *c* binding sites at CIII and CIV. Comparison of the integrated density for cyt. *c* in this map to the two other maps each with a well-defined single cyt. *c* bound at CIII or CIV indicates a cyt. *c* occupancy of 1.1:1:1, respectively. Therefore, despite its low resolution, the cyt. *c* density in the intermediate position is not consistent with multiple cyt. *c* molecules bound to the supercomplex at the same time. Instead, it indicates that in the intermediate position between CIII and CIV cyt. *c* retains positional variability that could not be resolved by the principal component analysis. This observation suggests that there is an approximately equal probability of cyt. *c* binding CIII or CIV, or occupying the stretch of supercomplex surface between these two sites.

Cytochrome *c* has a net positive charge and a dipole moment (44), with its heme *c* group positioned with its edge near the positively charged protein surface. The atomic models of cyt. *c* bound to CIII and CIV (42, 43) show that it receives and donates its electron with its positively-charged surface facing its binding sites. The cyt. *c* positions detected by cryo-EM lie along a negatively-charged path on the supercomplex surface (36, 45) (**Fig. 4F**). Together, these data indicate that supercomplex formation allows association of a single cyt. *c* with this negatively-charged path. Cytochrome *c* can then travel along this path while maintaining its positively-charged surface oriented toward the supercomplex surface, thereby allowing cyt. *c* to diffuse along the surface between its binding sites on CIII and CIV.

## Discussion

Together, the kinetic and structural data indicate that electron transfer between CIII and CIV in the supercomplex is mediated by 2D diffusion of a surface-associated cyt. *c*. This association is facilitated by a negatively charged path that increases the affinity of the positively charged cyt. *c* for the supercomplex (see also (33, 46)). While two of the three cyt. *c* positions observed by cryo-EM are consistent with earlier X-ray crystal structures (42, 43), the third position, intermediate between CIII and CIV, is unique to the CIII_2_CIV_1/2_ supercomplex. The nearly equal occupancy of the three positions presumably reflects the dynamics of the weakly interacting cyt. *c* that allow it to slide along the surface between the electron donor and acceptor sites. Thus, association of CIII_2_ and CIV to form a supercomplex does not enhance electron transfer by decreasing the distance cyt. *c*. diffuses in 3D (47), but rather enables a different mechanism for electron transfer with 2D diffusion of cyt. *c*.

Assuming the 2D diffusion of cyt. *c* between its binding sites at CIII and CIV, which are ~10 nm apart, occurs with a time constant of ~50 msec (~20 s^-1^, **Fig. 2A**) at cyt. *c*:supercomplex ratios of 1 to ~10, yields a 2D diffusion coefficient of 5×10^-12^ cm^2^/s (*L*^2^=4 *D t*). This value is approximately 100 smaller than the 2D diffusion coefficient obtained for cyt. *c* on a mitochondrial membrane surface (32, 48). The difference in coefficients suggests that the electrostatic interactions between cyt. *c* and the negatively charged supercomplex surface seen by cryo-EM are more specific than between cyt. *c* and the average mitochondrial membrane.

At cyt. *c*:supercomplex ratios of 1 to ~10, electron transfer between CIII and CIV involves a single associated cyt. *c*, as evident from the minimal increase in rate as the concentration of cyt. *c* is increased in this range (**Fig. 2A**). In contrast, the increase in supercomplex activity with cyt. *c*:supercomplex ratios above ~10 indicates involvement of additional cyt. *c* molecules at higher ratios. The additional cyt. *c* could either bind transiently at the supercomplex surface to establish a connection via two or more bound cyt. *c* molecules or mediate electron transfer via 3D diffusion. Unfortunately, cryo-EM of the supercomplex with high concentrations of cyt. *c* is precluded by background in images due to the excess of cyt. *c*. However, the decrease in supercomplex activity from ~60 e^-^/s to ~40 e^-^/s upon dissociation of the supercomplex in DDM (**Fig. 2C**) supports the former explanation. Electron transfer from CIII to CIV via two cyt. *c* molecules resembles the stable supercomplex from *M. smegmatis*, where a di-heme cyt. *cc* component of the supercomplex electronically connects CIII and CIV (49, 50). The concentration of cyt. *c* in the yeast intermembrane space was found previously to be 100 to 500 μM (8), with a cyt. *c*:CIV ratio of ~5 (31, 51). Therefore, *in vivo*, electron transfer from CIII to CIV would be expected to involve at least one cyt. *c* associated with the supercomplex surface.

The binding of cyt. *c* to the supercomplex is in accord with kinetic studies of *S. cerevisiae* mitochondria, which showed a tightly bound cyt. *c* for each CIV in the mitochondrial inner membrane (51), as well as steady-state kinetic studies of electron transfer between CIII and CIV (52). The observation is also consistent with the finding that cyt. *c* co-purifies with the mouse supercomplexes (14). Furthermore, multiple interactions between cyt. *c* and CIII detected by NMR and ITC have been suggested to facilitate electron transfer to CIV via “sliding” of cyt. *c* within a supercomplex (46).

The functional role of supercomplexes has been debated extensively in the literature (e.g. (12, 17, 34, 53–55)). As evident from the present study, changes in turnover activity of individual respiratory complexes upon forming supercomplexes are too small to result in functionally relevant changes in the electron flux through the respiratory chain (see also (34, 53)). Furthermore, the interaction surfaces of the supercomplex components in different organisms are highly variable, which suggests that association of the components does not result in specific modulation of function. Thus, the only common feature of supercomplexes is the decreased distance between their components.

The 2D diffusion of cyt. *c* resembles the “substrate channeling” model of respiratory chains, where cyt. *c* diffusion is restricted to the space surrounding the supercomplex. This model has been criticized based on the finding that cyt. *c* diffusion in *S. cerevisiae* is unrestricted (51). However, the scenario suggested by our data is that weak electrostatic interactions between the positively charged cyt. *c* and the negatively charged supercomplex surface lead to the surface-association of cyt. *c*, which remains in equilibrium with the cyt. *c* pool (8, 51).

In addition to differences in cyt. *c* diffusion rates, 2D and 3D diffusion of cyt. *c* between CIII and CIV would lead to differences in the redox state of the cyt. *c* pool. With 2D diffusion, the redox state of the bulk cyt. *c* pool is determined by the equilibrium constant for exchange between the surface-associated cyt. *c* and the bulk cyt. *c* pool. For 3D diffusion, the redox state of the bulk cyt. *c* depends only on the relative turnover rates of CIII and CIV. Consequently, formation of supercomplexes should perturb the reduced:oxidized cyt. *c* ratio of the pool. Because cyt. *c* is involved in an intricate web of redox interactions (56), regulated formation or dissociation of supercomplexes upon changing environmental conditions may result in triggering of cellular signaling pathways.

## Materials and Methods

### Strain and cell growth

The yeast strain BY 4741 from *S. cerevisiae* with a FLAG tag on CIV subunit Cox6 was used (35). Yeast cultures were grown in YPG (2% peptone, 1% yeast extract, 2% glycerol) at 30°C while shaking at 180 rpm.

### Preparation of mitochondrial membranes

Cells were harvested by centrifugation at 6500 × g (5 min, 4°C), washed with 50 mM potassium phosphate (KPi) buffer at pH 7, and then re-suspended in 650 mM Mannitol, 50 mM KPi buffer, 5 mM EDTA, pH 7.4. Cells were lysed by mechanical disruption (Constant Systems cell disrupter) at a pressure of 35 kpsi and cell debris was removed by centrifugation at 5500 × g (20 min, 4°C). The supernatant was then centrifuged at 120 000 × g for 1 h at 4°C. The pellet was resuspended and homogenized in 100 mM KCl, 50 mM KPi buffer, 5 mM EDTA, pH 7.4, centrifuged at 120 000 × g for 30 min at 4°C and washed two times with 50 mM KPi buffer, 5 mM EDTA, pH 7.4 and centrifuged as in the previous step. Finally, the pellet was homogenized in 50 mM KPi buffer, pH 7.4, frozen in liquid nitrogen, and stored at −80°C.

### Isolation of supercomplexes

Membrane fragments were diluted to a total protein concentration of 2 mg/ml in 100 mM KCl, 50 mM KPi buffer, pH 7.4, 0.5 % (w/v) glyco-diosgenin (GDN101, Anatrace) and solubilized overnight at 4°C. The membrane lysate was centrifuged at 140 000 × g for 1 h 30 min at 4°C. The cleared lysate was diluted to a final GDN concentration of 0.2 % and concentrated by centrifugation to roughly 70 ml with a 100 kDa molecular weight cut-off concentrator (Merck Millipore) before incubating with Anti-FLAG M2 resin (Sigma) for 2 h at 4°C. The Anti-FLAG M2 resin with protein bound was washed with 20 ml of wash buffer (150 mM KCl, 20 mM KPi buffer, pH 7.4, 0.01 % GDN) and eluted with 0.1 mg/ml FLAG peptide (Sigma), 150 mM KCl, 20 mM KPi buffer, pH 7.4, 0.01 % GDN. Eluted supercomplex was concentrated with a 100 kDa molecular weight cut-off concentrator (Merck Millipore) and further purified by size exclusion chromatography with an Äkta Pure M25 (GE Healthcare) operated at 4°C with UV detection at 280 nm and 415 nm. The sample was loaded on a Superose 6 Increase 10/300 GL column (GE Healthcare) equilibrated with 150 mM KCl, 20 mM KPi buffer, pH 7.4, 0.01 % GDN. Fractions of 250 μl were collected and fractions containing both CIII and CIV were pooled and concentrated as described above.

### UV-visible difference spectroscopy

Optical absorption spectra were recorded with a Cary 100 UV-VIS spectrophotometer (Agilent Technologies). A small aliquot of sodium dithionite was used as a reducing agent to obtain a reduced minus oxidized difference spectrum. Peaks were fitted for the different heme groups, heme *c*_1_ (554 nm), hemes *b*_H_ and *b*_L_ (562 nm), hemes *a* and *a*_3_ (603 nm) and their concentrations were calculated using the following difference absorption coefficients: ε_554_ = 21 mM^-1^cm^-1^ (57), ε_562_ = 51.2 mM^-1^cm^-1^ (see (58)) and ε_603_ = 25 mM^-1^cm^-1^ (59).

### Activity measurements

Decylubiquinone (Sigma) was dissolved in 100 % ethanol and reduced with several crystals of sodium borohydride (NaBH_4_, Sigma) to obtain decylubiquinol (DQH_2_). When the solution was clear, 10-20 μl 5 M HCl (depending on the amount of NaBH_4_ used for reduction) was added. The sample was then centrifuged for 10 min at 10000 × g and the supernatant containing DQH_2_ was collected.

The activity of CIII was measured by following absorbance changes as a function of time at 550 nm (reduction of cyt. *c*) with a Cary 100 UV-VIS spectrophotometer. A baseline absorbance was first recorded with 50 μM oxidized cyt. *c* from *S. cerevisiae* (Sigma), CIII_2_CIV_1/2_ supercomplex (equivalent of 20 nM CIIICIV), 2 mM KCN (to block cyt. *c* oxidation by CIV) in 150 mM KCl, 20 mM KPi buffer, pH 7.4, 0.01 % GDN. The reaction was initiated by addition of 100 μM DQH_2_ (from a 20 mM solution in ethanol). Control experiments showed that the reduction rate of cyt. *c* by DQH_2_ without supercomplex was negligible.

The activity of CIV was measured by monitoring the oxygen-reduction rate with a Clark-type oxygen electrode (Oxygraph, Hansatech) operated at 25°C. A baseline oxygen concentration was first recorded with 50 μM oxidized cyt. *c* from *S. cerevisiae*, 10 mM ascorbate, 100 μM N,N,N’,N’-Tetramethyl-p-phenylenediamine (TMPD) in 150 mM KCl, 20 mM KPi buffer, pH 7.4, and 0.01 % GDN. The reaction was initiated by addition of 10 nM supercomplex. The activity of CIV (10 nM) was also measured with CIV isolated from the CIV peak in **Fig. S1A** in either 0.01 % GDN or 0.05 % DDM.

To measure the activity of dissociated complexes CIII_2_ and CIV, DDM was added to a concentration of 0.5 % in a sample containing the supercomplex (CIII_2_CIV_1/2_) in GDN101. After incubation for 20 min the sample was diluted in the oxygraph chamber to yield a final concentration of 0.05 % DDM.

The QH_2_:O_2_ oxidoreductase activity of the supercomplex was measured by monitoring the oxygen-reduction rate using a Clark-type oxygen electrode operated at 25 °C. A baseline oxygen concentration was first recorded with an equivalent of 20 nM CIIICIV supercomplex, cyt. *c* from equine heart or *S. cerevisiae* (the cyt. *c* concentration was varied in the range 10 nM to 100 μM) in 150 mM KCl, 20 mM KPi buffer, pH 7.4, 0.01 % GDN. The reaction was initiated by addition of 100 μM DQH_2_.

### Gel electrophoresis

Blue native (BN) PAGE was performed with pre-cast gel, NativePAGE^™^ 4-16 % Bis-Tris (Invitrogen), according to the manufacturer’s instruction using Native PAGE^™^ 1X running buffer as anode buffer and NativePAGE™ 1X running buffer supplemented with NativePAGE^™^ 1X cathode buffer additive as cathode buffer (Invitrogen). The gel was run at 4 °C for 60 min at 150 V, before exchanging the cathode buffer to the anode buffer and running for an additional 40 min at 250 V. The gel was then stained with Coomassie Brilliant Blue.

### Grid preparation and cryo-electron microscopy

Purified supercomplexes (3 μl) at a concentration of 8 mg ml^-1^, supplemented with *S. cerevisiae* cyt. *c* (Sigma, aerobically grown *S. cerevisiae* contains 95 % isoform-1 cyt. *c* (60)) at a ~1:12 molar ratio, was applied to homemade nanofabricated holey gold grids (61–63) that had been glow-discharged in air (120 s, 20 mA using PELCO easiGlow). Grids were blotted for 3 s at 4 °C and 100 % humidity before rapid freezing in liquid ethane with a Vitrobot Mark IV (Thermo Fischer Scientific). Cryo-EM data were collected using a Titan Krios G3 electron microscope (Thermo Fisher Scientific) operated at 300 kV equipped with a prototype Falcon 4 direct detector device camera. Automated data collection was done with the EPU software package. An initial dataset of 5,690 movies, each consisting of 30 exposure fractions was collected at a nominal magnification of 75,000×, corresponding to a calibrated pixel size of 1.03 Å. The camera exposure rate and the total exposure of the specimen were 4.6 e^-^/pixel/s and ~42 e^-^/Å^2^, respectively (**Table S1**).

### Cryo-EM image processing

All image analysis was performed with cryoSPARC v2 (64). Movies were aligned with *MotionCor2* (65) and contrast transfer function (CTF) parameters were estimated in patches with a 7×7 grid. The dataset was manually curated to remove movies with devitrification, large cracks in the ice, or poor CTF fit parameters, reducing the dataset size to 5,147 movies. Templates for particle selection were generated by 2D classification of manually selected particles image leading to 423,969 selected particles. After particle selection, images were corrected for local motion (66) and extracted in 380×380 pixel boxes (**Table S1**). Extracted particles images were cleaned with six rounds of *ab initio* 3D classification and heterogeneous refinement, taking only the classes that corresponded to CIII_2_CIV_1_ and CIII_2_CIV_2_ after each round. This procedure further reduced the size of the dataset to 73,542 particle images with 49,273 and 24,269 particle images belonging to the CIII_2_CIV_1_ and CIII_2_CIV_2_ classes, respectively. These two classes were combined and used to refined a map of CIII_2_CIV_1_ with non-uniform refinement (67) and then analyzed with 3D variability analysis (3DVA) (68) with a mask over the region of the map corresponding to cyt. *c*. Cluster analysis was used to generate three structures that served as initial references in a heterogeneous refinement. The three populations resulting from the heterogeneous refinement corresponded to the CIII-bound, CIV-bound, and in-between populations (see **Fig. S6** for full workflow). The maps were locally filtered for presentation in figures. Particle images for the CIII-bound and CIV-bound populations were individually subject to particle subtraction and local non-uniform refinement with masks over the cyt. *c*-CIII_2_ and cyt. *c*-CIV regions, respectively. Rigid body fitting of cyt. *c* was performed with UCSF Chimera (69) based on existing structures (42, 43). Occupancy calculations were performed with custom Python programs (https://github.com/justinditrani). Maps were put on the same greyscale with diffmap (70) and occupancy was estimated by integrating the values of voxels within the mask used for 3DVA (**Fig. S4**) for the different cyt. *c* positions and dividing this value by the number of voxels in the mask.

## Data deposition

Data deposition: All electron cryomicroscopy maps described in this article have been deposited in the Electron Microscopy Data Bank (EMDB) (accession nos. EMD-XXXX to EMD-XXXX).

## Acknowledgement

We thank Irina Smirnova for experimental assistance in the initial phase of the project and Pia Ädelroth for valuable discussions. This work was supported by the Knut and Wallenberg Foundation (PB), the Swedish Research Council (PB), and the Canadian Institutes of Health Research grant PJT162186 (JLR). JDT was supported by a Canadian Institutes of Health Research Postdoctoral Fellowship and JLR was supported by the Canada Research Chairs program. Titan Krios cryo-EM data were collected at the Toronto High-Resolution High-Throughput cryo-EM facility supported by the Canada Foundation for Innovation and Ontario Research Fund. Molecular graphics and analyses performed with UCSF Chimera, developed by the Resource for Biocomputing, Visualization, and Informatics at the University of California, San Francisco, with support from NIH P41-GM103311.

## Author Contributions

P.B. and J.L.R. designed research; A.M. and J.D.T. performed research; J.D.T., A.M., J.L.R. and P.B. analyzed and interpreted the data; J.L.R. supervised the electron cryomicroscopy studies; P.B. supervised the kinetic studies; and J.L.R., P.B., A.M. and J.D.T. wrote the paper.

## Classification

Biological Sciences/Biochemistry.

