## Supplementary information for "Cryo-EM structure and kinetics reveal electron transfer by 2D diffusion of cytochrome *c* in the yeast III-IV respiratory supercomplex"

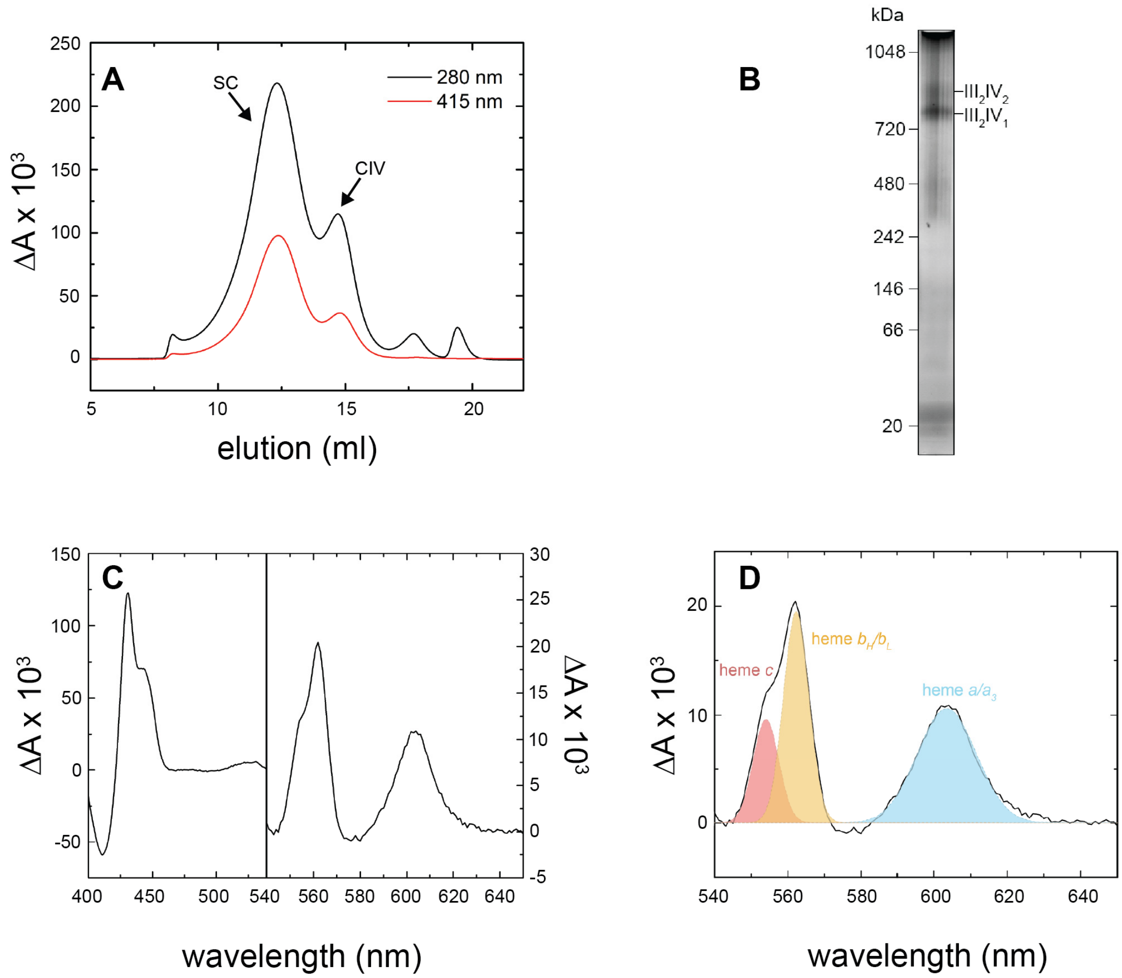


**Figure S1**. **Purification of the yeast Complex III/IV supercomplex.** (**A**) Elution profile from size-exclusion chromatography with a Superose 6 Increase 10/300 GL column in 150 mM KCl, 20 mM KPi buffer, pH 7.4, 0.01 % GDN. Elution was followed at 280 nm (total protein) and 415 nm (hemes). Hemes (415 nm) were found in two of the peaks. Dithionite reduced minus oxidized difference spectra of the fractions showed the presence of supercomplexes (SC), eluted in the first peak, and free CIV in the second peak. (**B**) BN-PAGE of the purified supercomplex (12-ml peak in panel A) showing presence of both III$_2$IV$_1$ and III$_2$IV$_2$ supercomplexes. (**C**) Dithionite reduced-minus-oxidized difference spectrum of purified supercomplex after size exclusion chromatography (see panel A) in 150 mM KCl, 20 mM KPi buffer, pH 7.4, 0.01% GDN. (**D**) Peaks from the difference spectrum in panel (C) were fitted using Origin Pro 2016 (OriginLab corporation). The following difference absorption coefficients were used: $\varepsilon_{554}$ = 21 mM$^{-1}$cm$^{-1}$ (1), $\varepsilon_{562}$ = 51.2 mM$^{-1}$cm$^{-1}$ (see (2)) and $\varepsilon_{603}$ = 25 mM$^{-1}$cm$^{-1}$ (3).

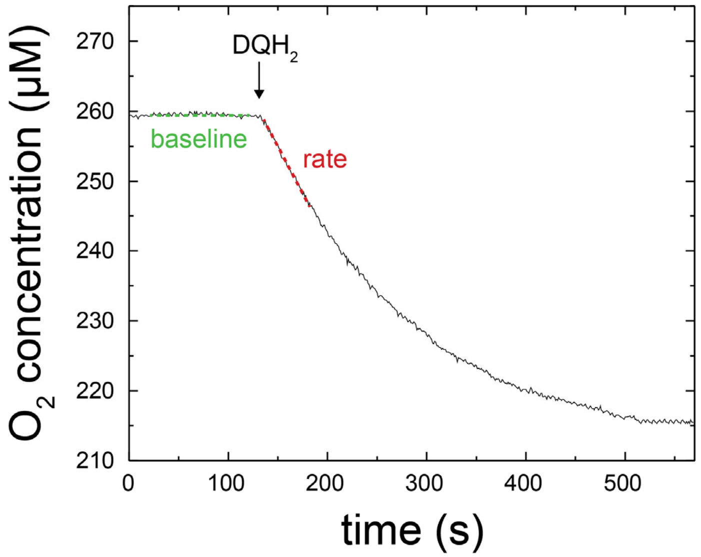


**Figure S2**. **Oxygen consumption by the yeast Complex III/IV supercomplex.** The quinol oxidation-$O_2$ reduction rate of the *S. cerevisiae* supercomplex, measured by following reduction of $O_2$ over time. The turnover rate was determined from the difference of the slopes after (rate) and before (baseline) addition of $DQH_2$. The slope with only DQH2 (no enzyme) was approximately zero. Experimental conditions: 50 µM cyt. *c*, 100 µM decylubiquinol ($DQH_2$), 150 mM KCl, 20 mM $KP_i$ buffer, pH 7.4, 0.01 % GDN.

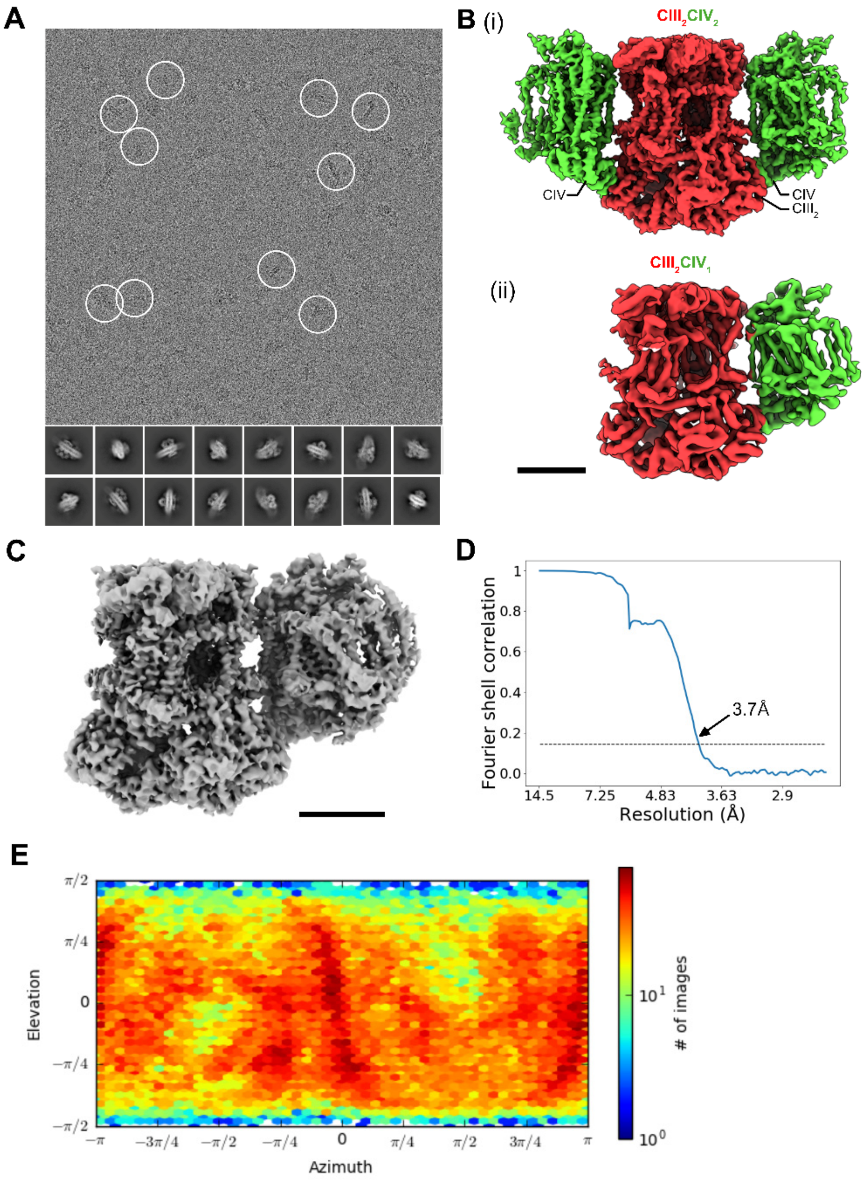


**Figure S3**. **CryoEM map calculation.** (**A**) Representative micrograph and 2D class averages for supercomplex. (**B**) CIII2CIV2 structure (***i***) and CIII2CIV structure (***ii***). (**C**) CIII2CIV map following local non-uniform refinement. (**D**) Fourier shell correlation (FSC) curve after correction for solvent masking. (**E**) Viewing direction distribution for particle images. Scale bars, 50 Å.

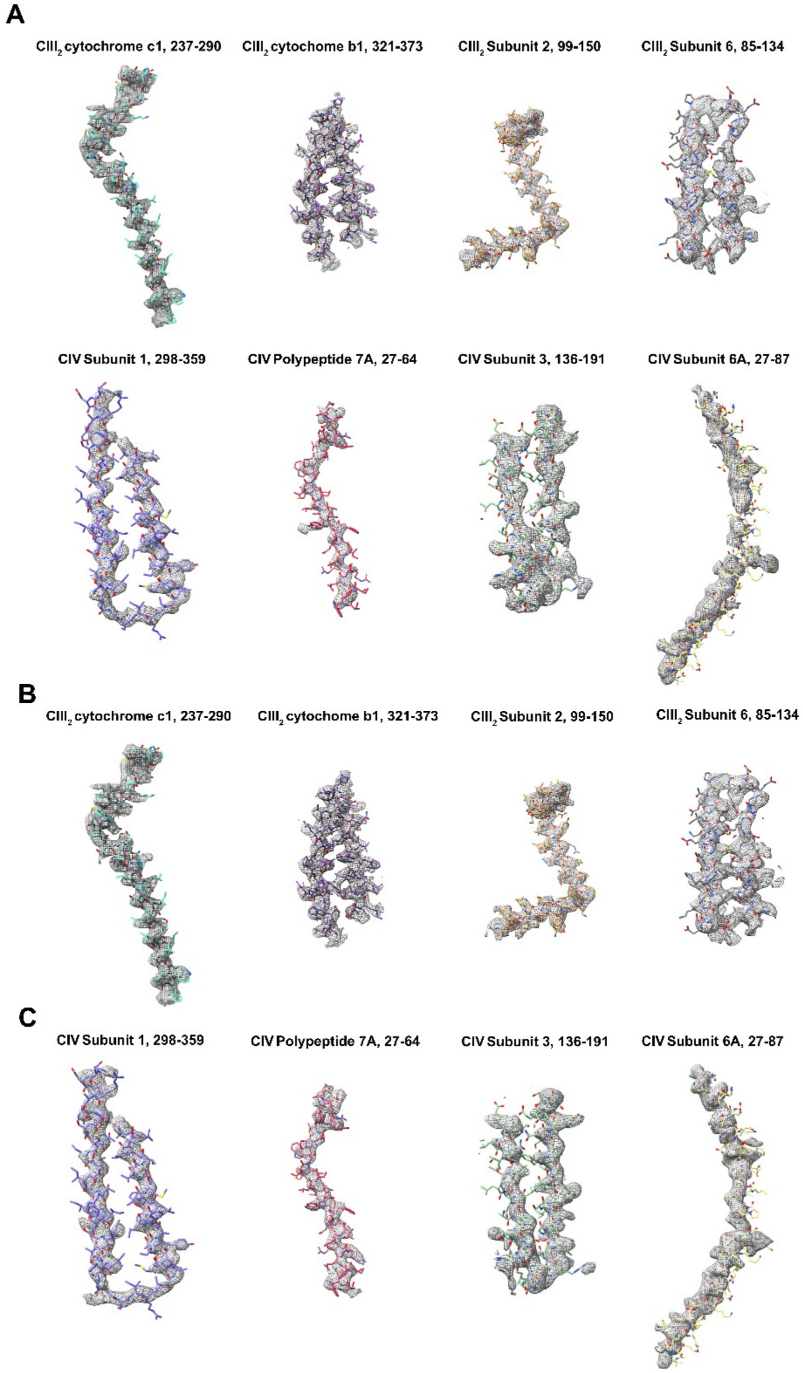


**Figure S4. Examples of model in map fit from cryoEM.** Examples are shown illustrating the variable resolution in the map including from $CIII_2CIV$ local refinement (**A**), $CIII_2$ local refinement (**B**), and CIV local refinement (**C**).

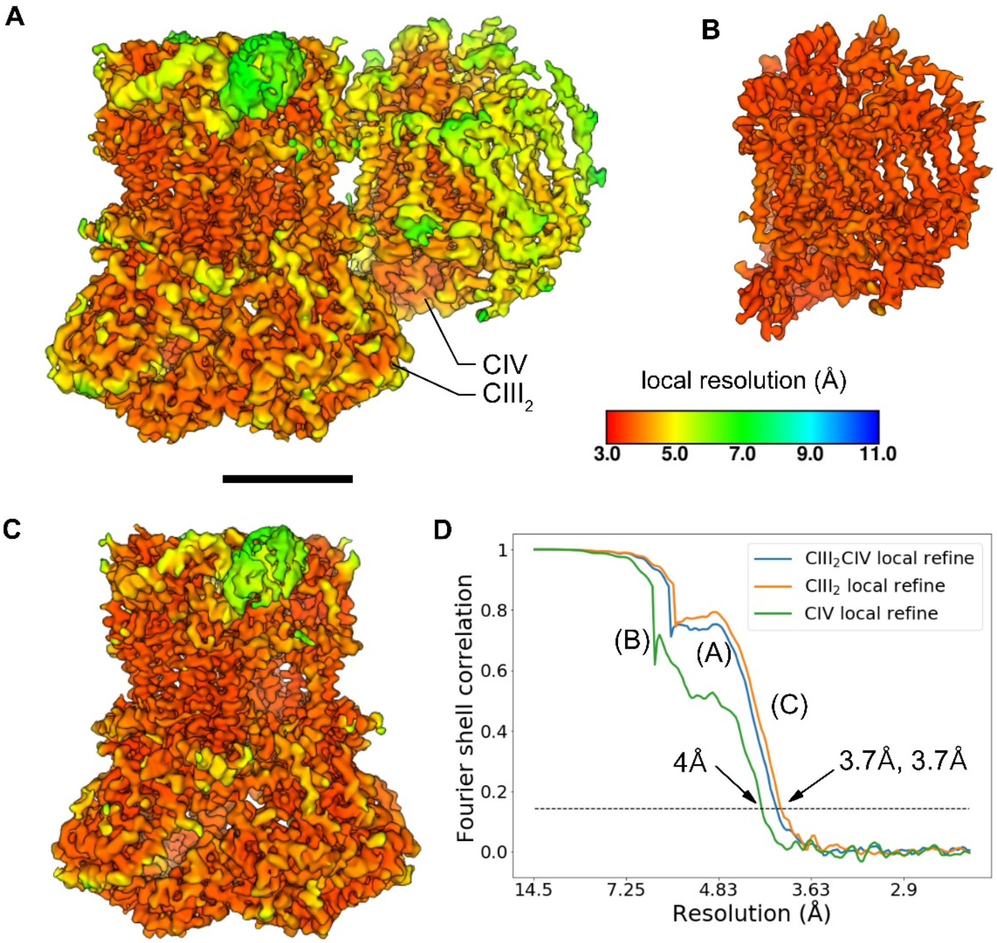


**Figure S5**. Local resolution estimates following local non-uniform refinement. (**A**) The $CIII_2CIV$ map. (**B**) CIV local refinement. (**C**) $CIII_2$ local refinement. (**D**) Overlay of Fourier shell correlation (FSC) curves corrected for solvent masking.

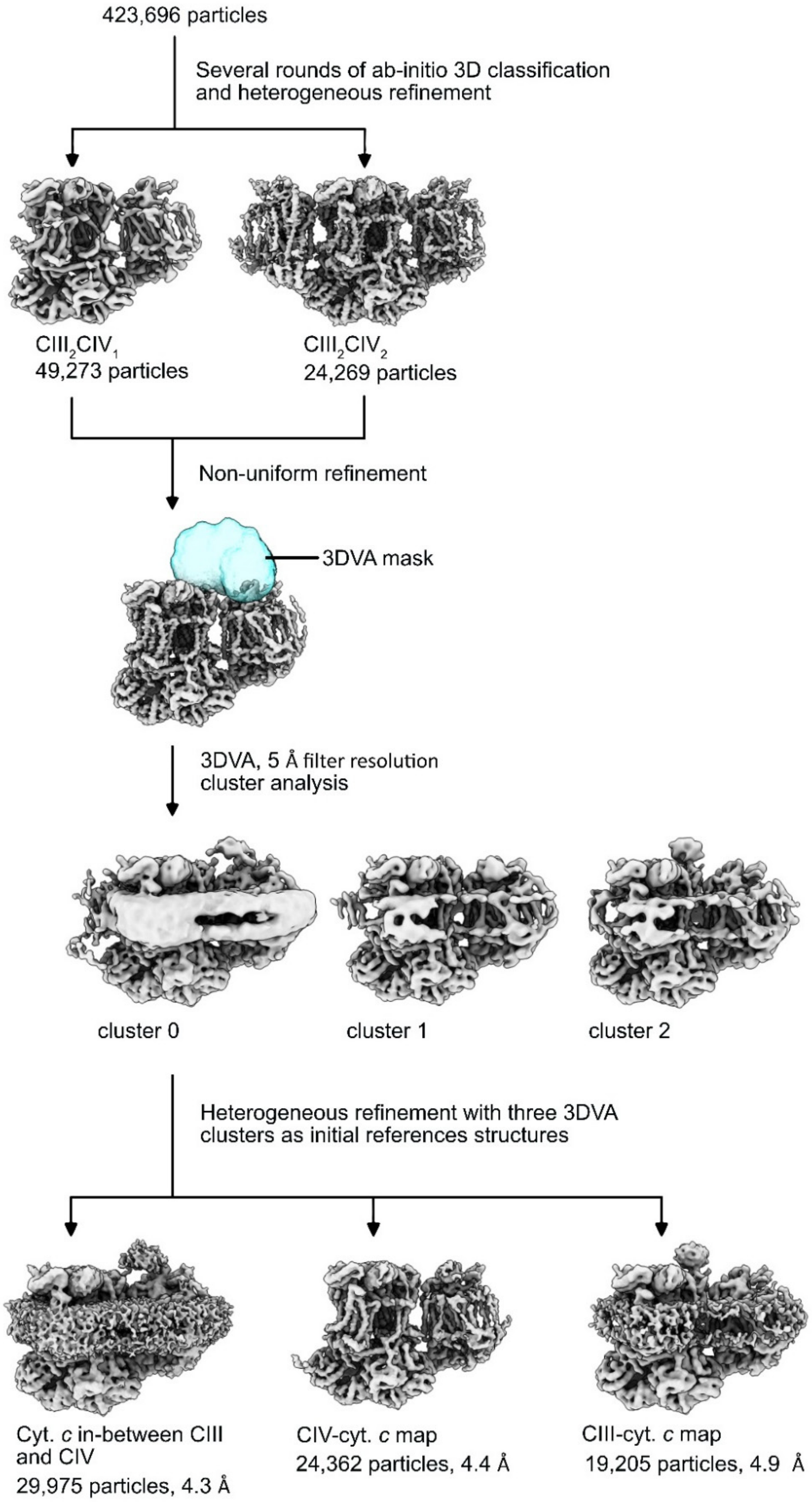


**Figure S6. Cryo-EM workflow.**

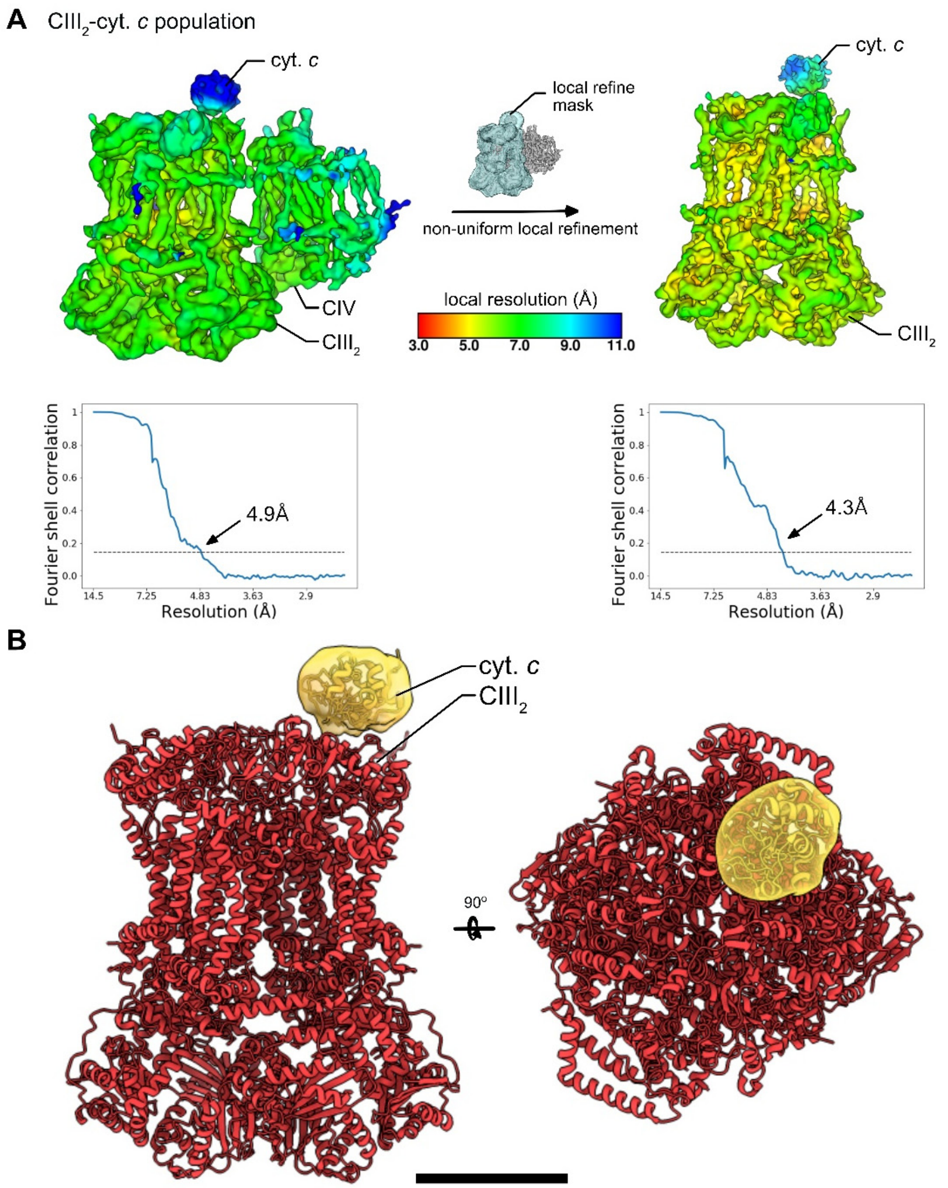


**Figure S7**. **Interaction of cyt. *c* with Complex $III_2$.** (**A**) Local resolution estimates and FSC curves corrected for solvent masking for $CIII_2$-cyt. *c* maps. (**B**) Side and top view of $CIII_2$ with cyt. *c* density (yellow) and the yeast cyt. *c* atomic model (PDB: 1YCC (4)) fit into the map. Scale bar, 50 Å

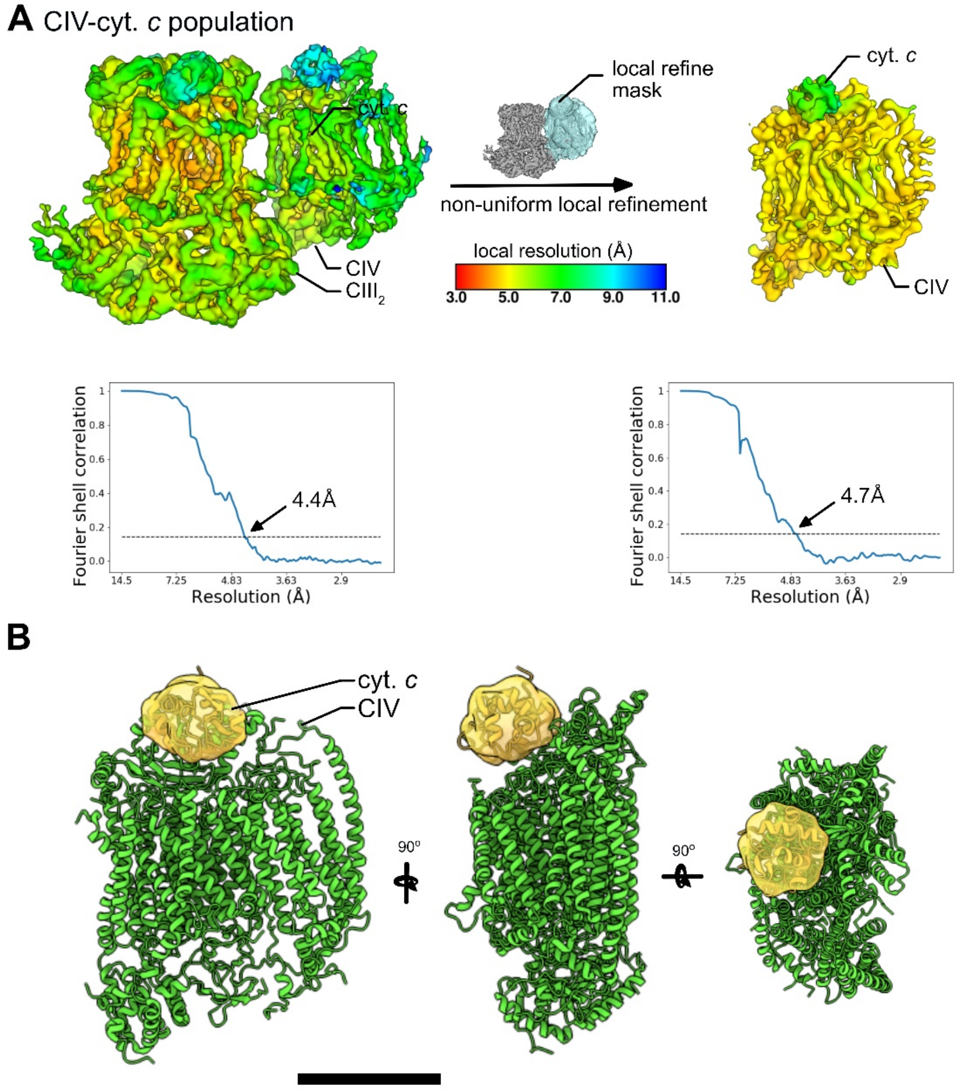


**Figure S8**. **Interaction of cyt. *c* with Complex IV.** (**A**) Local resolution estimates and FSC curves for CIV-cyt. *c* structures corrected for solvent masking. (**B**) Side and top view of $CIII_2$ with cyt *c*. density and yeast cyt *c*. atomic model (PDB: 1YCC (4)) fit into density. Scale bar, 50 Å.

**Table S1.** Cryo-EM data acquisition and image processing.

| **Data Collection** | |
|---|---|
| Electron Microscope | Titan Krios |
| Camera | Falcon 4 |
| Voltage (kV) | 300 |
| Nominal Magnification | 75,000 |
| Calibrated physical pixel size (Å) | 1.03 |
| Total exposure (e/Å$^2$) | 42 |
| Exposure rate (e/pixel/s) | 4.6 |
| Number of frames | 29 |
| Defocus range (μm) | 0.9 to 2 |
| **Image Processing** | |
| Motion correction software | *MotionCor2* |
| CTF estimation software | *cryoSPARC v2* |
| Particle selection software | *cryoSPARC v2* |
| Micrographs used | 5,690 |
| Particle images selected | 423,969 |
| 3D map classification and refinement software | *cryoSPARC v2* |
